# Mice expressing A53T / A30P mutant alpha-synuclein in dopamine neurons do not display behavioral deficits

**DOI:** 10.1101/2023.05.25.542172

**Authors:** Cameron Keomanivong, Josephine Schamp, Ervina Tabakovic, Ramasamy Thangavel, Georgina Aldridge, Andrew A. Pieper, Nandakumar S. Narayanan

**Author notes:** **Corresponding Author** Nandakumar Narayanan, 169 Newton Road, Pappajohn Biomedical Discovery Building—5336, University of Iowa, Iowa City, 52242. These authors contributed equally to this manuscript (Cameron Keomanivong, Josephine Schamp).

## Abstract

Alpha-synuclein has been implicated in neurodegenerative diseases such as Parkinson’s disease and Dementia with Lewy bodies, with A53T and A30P mutations shown to be disease-causing. It has been reported that transgenic mice with tyrosine hydroxylase promotor-driven expression of A53T / A30P mutant alpha-synuclein in dopamine neurons provide a useful preclinical model of these conditions by virtue of developing dopaminergic neuronal cell death and related behavioral deficits. Here, we report a lack of replication of this finding. Despite detecting robust overexpression of A53T / A30P mutant alpha-synuclein in dopamine neurons, we observed neither cell death or related behavioral deficits in these mice. Our results demonstrate that preclinical models of synucleinopathy need careful validation in the field.

## INTRODUCTION

Synucleinopathies such as Parkinson’s disease (PD) and Dementia with Lewy bodies (DLB) are poorly understood and difficult to treat, devastating neurodegenerative disorders. Alpha-synuclein is the major protein found in the large intraneuronal inclusions known as Lewy Bodies in PD and DLB patients (Spillantini et al., 1998). Laboratory assays of alpha-synuclein aggregation are abnormal in PD and DLB patients, and people with mutations or triplications in alpha-synuclein are at increased risk for developing PD or DLB (Singleton et al., 2003). Two common mutations associated with these conditions are A53T and A30P (Giasson et al., 2002).

To determine how alpha-synuclein contributes to neurodegeneration in these conditions, considerable effort has been devoted in the field to developing mouse models of synucleinopathy. This problem is complex because mice do not live as long as humans (∼2 years vs. 60+ years, respectively), do not have elaborated forebrain circuits like humans, and do not express the same forms of alpha-synuclein that are found in humans. In addition, mice without alpha-synuclein have only mild deficits (Abeliovich et al., 2000; Kokhan et al., 2012), and mice over-expressing alpha-synuclein through transgenic or viral techniques also have only mild behavioral deficits (Chesselet et al., 2012; Decressac et al., 2012; Kim et al., 2017). Notably, mice inoculated with alpha-synuclein protofibrils (PFFs) in the striatum have reliable behavioral deficits, while mice inoculated with the same PFFs in the cortex do not (Luk et al., 2012; Zhang et al., 2021). Prominent behavioral deficits have been reported in mice with globally expressed A53T mutant alpha-synuclein (Giasson et al., 2022), and mice with global expression of A53T / A30P mutant alpha-synuclein display dopaminergic cell loss and behavioral impairment (Kilpeläinen et al., 2019; Yan et al., 2017).

Expression of A53T / A30P mutant alpha-synuclein in dopaminergic neurons, under selective control of the tyrosine hydroxylase promoter, has been reported to specifically induce dopaminergic neuronal cell loss starting at 7-9 months of age, with behavioral deficits in manifesting from 13-23-month-months of age (Richfield et al., 2002). We attempted to utilize this reported preclinical model in our laboratory. However, after driving robust expression of A53T / A30P mutant alpha-synuclein in dopaminergic neurons and aging mice for 18 months, we did not observe any dopaminergic neuronal cell death or behavioral deficits. These data demonstrate that alpha-synuclein mouse models need to be carefully validated in the field to enable investigators to appropriately direct resources toward understanding and finding new ways to understand and treat neurodegenerative disease.

## METHODS

### Mice

Twenty hemizygous (10 female; C57BL/6J-Tg(Th-SNCA*A30P*A53T)39Eric/J mice were obtained from Jackson labs (hm^2^α-SYN-39), as well as 10 wild type littermate controls from the same litter (5 female). Mice were aged for 18 months prior to final behavioral assessments and histology. One control animal died during the experiment and was excluded. We completed experiments with 29 mice.

### Open-field behavior

Mice were placed in a clean open field container (40 × 40 cm) in a quiet, well-lit room. Motor and exploratory activity was video recorded and tracked for 10 min using ANY-maze software (Wood Dale, IL 60191). Total distance traveled was calculated by ANY-maze software. Odors were removed with 70% alcohol and the apparatus was allowed to air dry between each trial. Mice were evaluated in this assay at 3, 6, 12, and 15 months.

### Accelerating Rotarod

Mice underwent three consecutive days of training in which they were placed on a motorized rotarod (Rotamex) that increased rotation speed from 4 to 40 rpm over a 5-minute period. For each training day, mice were provided three opportunities to maintain their perch on the rotarod as rotational speed increased. On the third day, the time it took the mice to fall from the beam was recorded. Mice were evaluated in the rotarod assay at 3, 6, 12, and 16 months. At each age, mice were placed on the apparatus until they fell (for a maximum of 10 minutes). Each test week consisted of two days of testing, and mice were given 3 trials each day.

### Histology

At 18 months when experiments were complete, mice were euthanized by injections of 100 mg/kg ketamine and transcardially perfused with ice-cold PBS. Half the brain was placed in 4% paraformaldehyde, and half the brain was frozen for use in western blotting. The brains that were removed and post-fixed in paraformaldehyde overnight were immersed in 30% sucrose until the brains sank. Sections (40 μm) were made on a cryostat (Leica) and stored at 4 °C. For immunohistochemistry, free-floating sections were washed with PBS and then PBST (0.03% Triton-X100 in PBS), followed by incubation in 2% normal goat serum for 1 h at room temperature. Next, sections were incubated in primary antibody at 4 °C overnight. After washing with PBST, sections were incubated with secondary antibody for 1 h. After washing again with PBST, and then PBS, sections were mounted onto slides with mounting media containing DAPI (Invitrogen P36962). Sections were stained for tyrosine hydroxylase (Millpore-AB152) and synuclein (abcam ab27766). Images were captured on an Olympus VS120 Microscope or a Zeiss Confocal microscope. All fluorescent immunohistochemistry images were pseudocolored to represent multiple fluorophores (Figure 1A-D).

**Figure 1:**
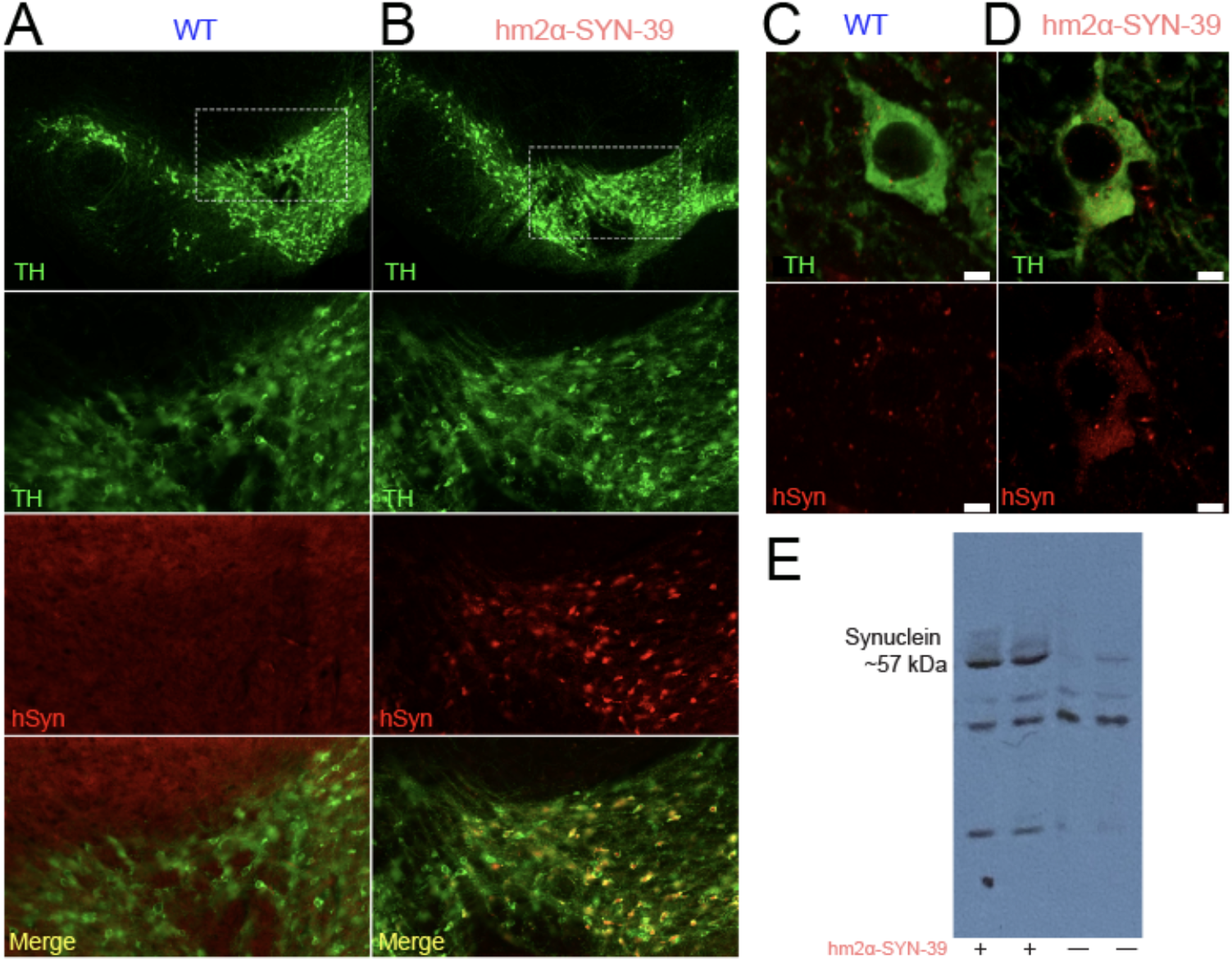
Low-power synuclein expression in an A) wild-type and B) A53T/A30P transgenic mouse. Top row; a coronal midbrain section through the substantia nigra stained for Tyrosine Hydroxylase (TH) in green; second row, a subsection of the white outline in the top row stained for TH in green. Third row; the same subsection stained for human synuclein in red; bottom row; a merged image with co-stained neurons in yellow. High-power images of synuclein expression in a C) wild-type and D) A53T/A30P transgenic mouse; scale bar = 5 μm. E) Western Blots from 2 A53T/A30P transgenic mice and two wild-type littermate controls.

### Western blotting

Protein concentrations per brain sample were determined utilizing a Pierce BCA protein assay kit (ThermoFisher). 50 ug of protein were loaded into each well and resolved by gel electrophoresis. Separated proteins were transferred from the gel to a nitrocellulose membrane, which was verified via Ponceau S staining. Membranes were blocked in TBST (0.1% Tween20 in TBS) with 5% BSA for 2 hours and then incubated with alpha-synuclein monoclonal primary antibody (LB509) overnight at 4°C. After washing with TBST, membranes were incubated with secondary antibody for 2 hours at room temperature. Membranes were washed again and then used for enhanced chemiluminescence utilizing the Bio-Rad Clarity Western ECL Substrate (Figure 1E).

## RESULTS

We tested the hypothesis that overexpression of alpha-synuclein in dopamine neurons would lead to neuronal loss and behavioral deficits by comparing 20 mice expressing A53T/A30P mutant alpha-synuclein under the tyrosine hydroxylase (TH) promotor (TH+; hm^2^α-SYN-39 mice) to 9 littermate wild-type mice. Overexpression of mutant alpha-synuclein was verified by immunohistochemistry, showing colocalization of alpha-synuclein with tyrosine hydroxylase (TH)-positive cells in the substantia nigra (Fig 1 A-B). Qualitative, blinded examination by two investigators confirmed increased expression of mutant human alpha-synuclein in the soma of TH+ neurons in the hm^2^α-SYN-39 model compared with wildtype controls by confocal microscopy (Fig 1 C-D). Western blotting also confirmed expression of mutant alpha-synuclein in transgenic mice (Fig 1E), whereas wild-type littermate mice did not express human alpha-synuclein. Notably, we did not observe any reduction in dopaminergic neuronal cells in transgenic mice compared to wild-type littermates (Fig 1A-B).

Next, we evaluated for behavioral deficits through testing in the accelerating rotarod and open-field tasks, in which it was previously reported that these mice displayed deficits at 15 months of age. Performance in the accelerating rotarod task was quantified by measuring the time mice stayed on the rotarod (Fig 2A) and the maximum revolutions per minute attained at the point of falling (Fig 2B). For rotarod performance over 3 – 16 months, linear-mixed-effects models revealed no main effects of time, synucleinopathy, or higher interactions. In the open field test, we monitored distance traveled (Fig 2C). Though there was a significant effect of time (F=21.8; p<0.00005), an effect of genotype (F=4.4; p<0.04), and an interaction of time with genotype (F=4.7; p<0.03), this was only because hm^2^α-SYN-39 mice were less active initially and then became hyperactive with age (Fig 2C). Importantly, at 15 months of age, there was no difference in performance between transgenic mice and wild-type littermates, in contrast to what was previously reported (Richfield et al., 2022). Thus, the previous report of behavioral deficits in hm^2^α-SYN-39 mice expressing A53T/A30P mutations cannot be replicated.

**Figure 2:**
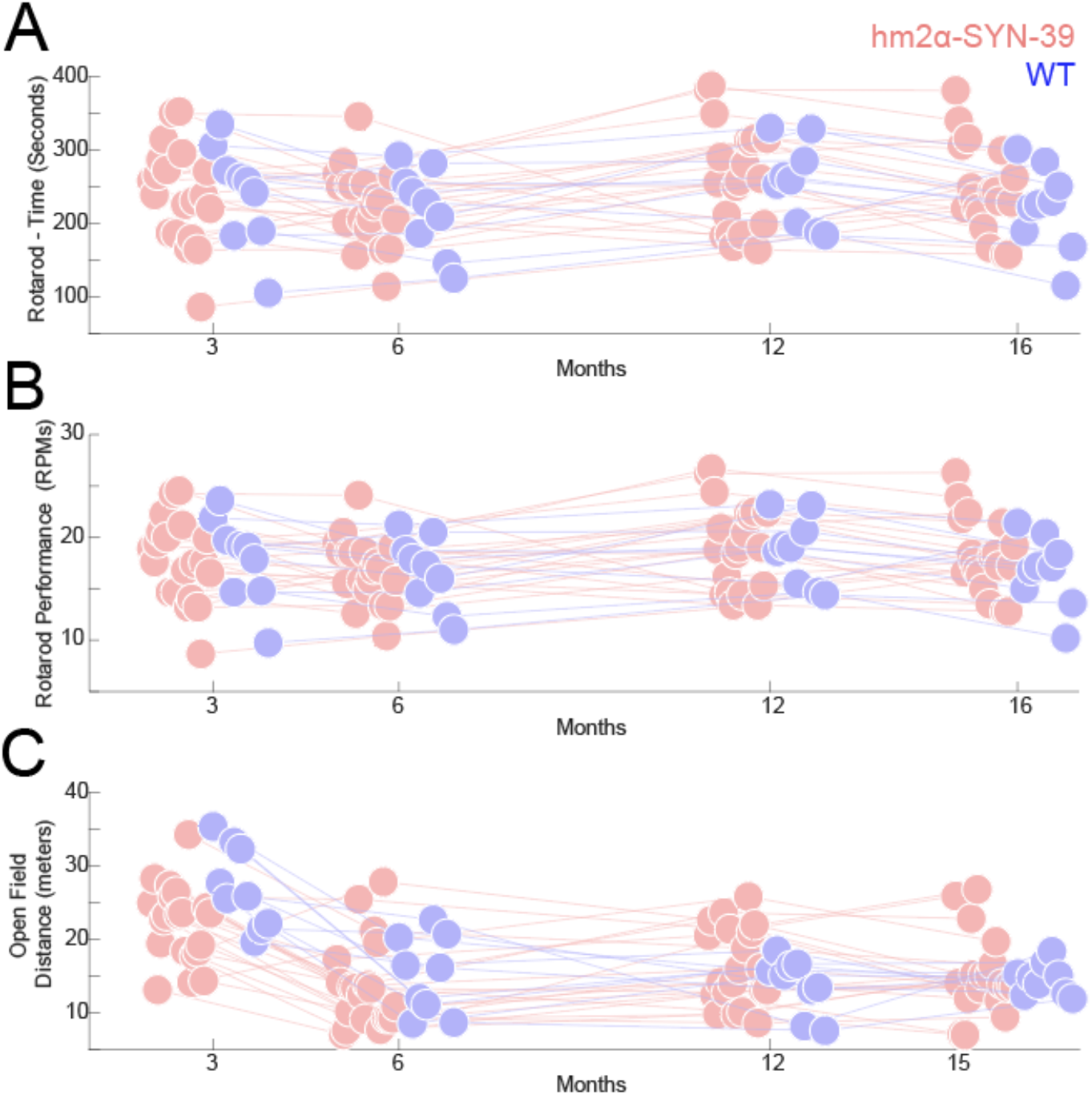
A) Time on the rotarod and B) maximum revolutions-per-minute. There were no reliable effects of genotype for 20 hm^2^α-SYN-39 mice (red) vs 9 wild-type mice (WT; blue). C) Open-field distance traveled for hm^2^α-SYN-39 mice vs. WT mice; there was a main effect of time, genotype, and an interaction due to the hm^2^α-SYN-39 mice being less active early and hyperactive late in their lives. Points were jittered for visualization.

## DISCUSSION

We investigated if mice expressing the A53T/A30P mutation in alpha-synuclein in dopaminergic neurons had neurodegeneration and behavioral deficits and found neither to be the case. We found no evidence of behavioral deficits in rotarod or open-field behavior, in contrast with the prior report (Richfield et al. 2002). These data suggest that this alpha-synuclein model does not reliably produce neurodegeneration or behavioral deficits, suggesting that alpha-synuclein models must be carefully validated. There were differences in open-field behavior as a function of genotype, but these mice were less active initially and hyperactive when aged, making this an inconsistent finding. Furthermore, these behavioral findings are opposite those noted in published literature, which reported hypoactivity at later ages (Richfield et al., 2002).

The report of this finding in these mice was published nearly 20 years ago, and there may have been substantial genetic drift or changes since then. A recent study further bred this mouse line to be homozygous with additional copies of the A53T/A30P mutation and found behavioral deficits at 3 months (Kilpeläinen et al., 2019). In addition, this study used only male mice, whereas we included males and females. Also, there may be distinct environmental differences that affect synuclein pathology, such as variances in the microbiome or other aspects of inflammation (Berardi et al., 2007). Lastly, publication bias in the field may have resulted in other laboratories also having found negative findings with this mouse line but not publishing their results. Our hope is that publication of our results will add clarity and direction for the field of synuclein-related neurodegeneration.

## CONTRIBUTIONS

AAP, JS, and NSN designed the experiments. CK and JS conducted the experiments. CK, JS, ET, RT, and GA analyzed data. AAP, GMA, and NSN wrote the paper

